# From Snapshots to Structure: A Novel Method for Reconstructing Directed Microbial Interaction Networks from Compositional Data

**DOI:** 10.1101/2025.08.16.670688

**Authors:** N Ranjith

## Abstract

Microbial communities are shaped by complex ecological interactions, but inferring these from 16S rRNA gene sequencing remains challenging due to data compositionality and the limitations of correlation-based methods. We present a novel framework that reconstructs directed, signed, and weighted microbial interaction networks from cross-sectional compositional data, without requiring time-series or predefined dynamic models. Using asymmetric slopes and a perturbation-informed strategy, This method infers interaction polarity and strength while accounting for compositional constraints. Applied to a synthetic gut microbiome, the framework uncovered taxon-specific trajectories, directional dependencies, and persistent interaction plasticity. Persistent Directed Acyclic Graph motifs identified keystone initiators, while other taxa served as resilient hubs. A quadrant-based visualization clarified ecological influencers and responders. Inferred networks aligned interaction polarity with abundance trends, revealing a principle of polarity-driven succession. This approach enables predictive, mechanistic insights into microbial dynamics from static data, offering scalable tools for microbiome analysis, synthetic community design, and ecological theory development.

## 1 INTRODUCTION

Microbial communities undergo continual restructuring in response to environmental change, internal competition, and resource availability. These shifts play a foundational role in ecosystem functioning, influencing processes such as nutrient cycling, host health, and community stability and plasticity. Yet the mechanisms governing how microbial species interact and co-assemble into stable or shifting communities remain poorly resolved, particularly under natural or semi-natural conditions. Network-based approaches have become central to studying these interactions, offering a means to capture ecological relationships such as mutualism, competition, and commensalism as edges between taxa. The structure of such networks can reveal influential species, emergent properties like resilience or modularity, plasticity and the pathways through which ecological effects propagate. However, most microbial networks inferred from 16S rRNA gene amplicon data rely on correlation or co-occurrence based methods, which yield undirected associations and do not capture causal directionality, interaction sign (positive or negative), or magnitude. As a result, they often produce visually dense and mechanistically ambiguous hairball networks that obscure ecological interpretation (Röttjers and Faust, 2018; Dohlman and Shen, 2019).

A second, and more fundamental, challenge arises from the compositional nature of amplicon sequencing data, where abundances are expressed as relative rather than absolute counts. This introduces statistical distortions unless properly corrected, typically via log-ratio transformations such as the centered log-ratio (CLR) (Gloor et al., 2017). While existing tools like SPIEC-EASI and SparCC attempt to address compositionality, they still produce undirected networks and are limited in ecological interpretability (Kurtz et al., 2015). Efforts to infer causal or directional interactions have turned to time-series and multi-omic datasets. While promising in theory, these methods face practical limitations. First, they typically rely on the selection of parameterized population dynamics models, which is challenging given the diversity of microbial interactions and nonlinear behaviors that can arise even in small systems (Xiao et al., 2017). Second, informative time-series data are difficult to obtain in many microbial systems, particularly host-associated communities such as the human gut microbiota, which often exhibit stability and resilience. As a result, available data largely reflect steady-state behavior, limiting the informativeness of dynamic models. Invasive perturbations aimed at increasing temporal variation may be impractical or ethically untenable in human or animal studies. Despite growing recognition of the dynamic nature of microbiome ecosystems, the mechanisms by which individual microbial populations contribute to and are shaped by ecological interactions remain poorly characterized (Kodera et al., 2022; Bernabe et al., 2018). This leaves a major methodological gap: how can we infer ecologically meaningful, directional interactions of individuals from static, compositional datasets which exhibits estabilishments of plasticity across time or conditions ?

To address this, a reproducible pipeline named DENIM (Directed Eco-logical Networks in Microbiomes) is introduced, which is designed to extract signed, directed, and weighted ecological interactions directly from CLR-transformed cross-sectional 16S data. This approach estimates asymmetric slopes between taxa to infer directionality and applies adaptive perturbation analysis to evaluate interaction sign and strength. In doing so, it avoids reliance on pre-specified dynamic models or longitudinal data, enabling mechanistic network inference from steady-state observations. The resulting networks are visualized using a biologically optimized layout the Eco-Cartesian framework which reduces visual clutter and highlights patterns of influence and response among taxa. Further consistent Directed Acyclic Graphs (DAGs) were identified across conditions, representing conditionally persistent interaction motifs that offer insight into temporally (conditional) established ecological interaction and higher-order ecological structure. These motifs capture microbial interaction plasticity (Solowiej-Wedderburn et al., 2025), hierarchical dependencies and directional pathways of influence that are otherwise masked in conventional network reconstructions.

By enabling the inference of interpretable, directional microbial interaction networks from widely available compositional datasets, this framework opens new avenues for microbial ecology. It supports predictive insights into community dynamics, enhances hypothesis generation for experimental testing of interaction plasticity, and provides a scalable tool for leveraging the growing troves of metagenomic and preexisting archived microbiome data collected across diverse environments (Kelliher et al., 2025; Oña et al., 2025).

Application of the method reveals a robust ecological principle, termed polarity-driven succession, in which the net balance of cooperative and antagonistic interactions predicts species abundance trajectories and overall community dynamics. This mechanistic insight advances microbial ecology by enabling predictive interpretations from static, cross-sectional datasets and broadening the theoretical understanding of microbial interaction networks. The results also support the hypothesis that pairwise linear associative interactions, when organized into a directed acyclic graph (DAG), can approximate nonlinear abundance dynamics by propagating cumulative net effects throughout the network.

## 2 METHOD

This pipeline is designed to reconstruct approximate ecological networks that capture not only associations but also the direction and type of species interactions, using only 16S rRNA gene amplicon data with replicate sampling under consistent conditions. Microbiome datasets are known to be compositional, meaning that the relative nature of the data can distort correlation and regression-based analyses. To mitigate these artifacts, the centered log-ratio (CLR) transformation is applied first (1), which appropriately normalizes abundance count data by adjusting each taxon’s abundance relative to the geometric mean of all taxa in the sample. This produces a transformed abundance matrix that better reflects underlying biological patterns. Following transformation, co-occurring taxa pairs were screened by retaining only those species that were simultaneously present in multiple replicates (at least three), thereby reducing biases from zero inflation. For each pair, both Pearson and Spearman correlations were computed across the datasets to gain additional insights. Pairs showing statistically significant association in either metric (at p < 0.05) were considered significant (Box 1).

Ecological interactions are rarely symmetric. For instance, one microbial species may inhibit another without being influenced in return (Kodera et al., 2022; Xiao et al., 2017). To capture such asymmetries, DENIM infers directionality by comparing the strength of pairwise linear regression slopes between taxa. Specifically, for each given taxon pairs (*x* and *y*), the summary statistics (2) along with their individual variability and their covariation were calculated (3) to understand how how strongly the abundance of the two taxa are related (covariance), and how much each taxon’s abundance varies on its own (variance). Using these statistics, the linear relationship in both directions, from taxon *x* to taxon *y*, and vice versa by fitting simple linear regression is calculated (4), where the slope of regression reflects the strength of the predictive relationship in that direction. Then by comparing the slope magnitudes of *y* regressed on *x* against the slope of *x* regressed on *y*, the dominant directional edge was inferred. If one slope is substantially larger (by a factor of 1.5 or more), this asymmetry is interpreted as evidence for a directed influence (5) (Blöbaum et al., 2019). This approach reflects the ecological intuition that if changes in the abundance of species *x* consistently predict changes in *y*, but not vice versa, *x* is more likely exerting a directional effect on *y*. Unlike temporal modeling or perturbation experiments, this method allows directional inference from steady-state cross-sectional data, a format that is highly common in microbiome studies.

After establishing directional links between taxa, The qualitative nature of these interactions are determined, whether they were likely to be positive (facilitative), negative (inhibitory), or neutral. To do this, a simple but ecologically intuitive approach is applied called local perturbation analysis (Box 2) (Medeiros et al., 2023). The central idea is to simulate small changes in the abundance of a given “source” taxon and observe how these changes would hypothetically affect its “target” partner, based on the slope of their statistical relationship. This mimics the concept of ecological sensitivity analysis, where minor changes in one species are used to estimate their downstream influence on another. For each directed pair (e.g., taxon *x* → taxon *y*), Where a small artificial perturbations is introduced to *x*’s abundance, both positively and negatively (6). These perturbations were scaled to be proportional to the original abundance specifically, 10% of the absolute value of each observation is used, with a minimum change threshold to ensure numerical stability (7). This adaptive scaling ensures that both rare and abundant taxa are perturbed in a biologically meaningful way. Using the previously estimated linear relationship between *x* and *y*, then the expected response in *y*’s abundance under each perturbed condition is calculated. A positive response to a positive perturbation suggests a facilitative (or cooperative) effect, while a negative response implies inhibition or competition. If both positive and negative perturbations yield the same directional response in *y* (e.g., both lead to an increase), this indicates potential non-linearity or context-dependent behavior if the slope is significantly less and is marked as ambiguous (8). All the inferred signs for every replicates in both +ve and −ve are collected and most frequent occurring sign is assigned for the corresponding directed interaction.

By repeating this process across all replicates and aggregating the results, each interaction is ultimately classified as positive, negative, neutral, or undetermined. This classification scheme provides a simple but informative annotation of the network’s edges which aids in reconstruction of polarity based approximate networks that explains species succession patterns, allowing for ecological interpretation of not just who affects whom, but how.

### 2.1 Network Construction and Visualization

To enhance the interpretability of microbial interaction networks, a novel visualization framework based on a Cartesian grid system was developed (Dohlman and Shen, 2019). In this approach, species were placed along the x and y axes according to their degree centrality within the network, replacing traditional numeric units with species names. Ecological interactions were then plotted (Fig 2) as directional edges within the appropriate quadrants of the grid, corresponding to their qualitative effects: mutualism (`++`), competition (`−−`), parasitism/predation (`+−` and `−+`) (Culp and Goodman, 2023; Dohlman and Shen, 2019; Oña et al., 2025). The magnitude of each interaction was represented by the size of slope-indicative bubbles. This design avoids node overlap and visual clutter commonly encountered in traditional network diagrams, providing a clearer representation of directional influence and interaction strength in simplified synthetic microbiome systems.

### 2.2 Cross-Condition DAG Construction and Comparison

Directed acyclic graphs (DAGs) are a well established framework in causal inference, forming a foundational component of path analysis and network cascade modeling. While DAGs are widely used in fields such as statistics and epidemiology, their application remains relatively limited in the biological sciences, particularly in microbial ecology (Borger and Ramesh, 2025). In this method, DAGs were used to represent directional dependencies and non-linear interactions within microbial taxa across multiple experimental conditions. To identify consistent (plastic) interactions, DAGs inferred from each condition-specific dataset were aggregated and compared. Recurrent subgraphs particularly stable three node motifs were identified by aligning DAG structures across conditions. These were subsequently visualized to highlight plastic interaction patterns that persist regardless of environmental variation or temporal stratification.

### 2.3 Analysis of empirical bacterial time series data

The DENIM was applied to an empirical time series dataset derived from a synthetic microbial community composed of 10 well characterized core human gut microbiome species (Shetty et al., 2022), making it an ideal model for inter-species ecological interaction analysis. This dataset was derived from 16s rRNA gene amplicon sequencing data, that comprises 20 samples: four biological replicates (B1–B4) sampled at five time points (T0, T24, T48, T72, and T96 hours). Raw OTU tables were obtained from the study’s GitHub repository. No additional taxonomic classification or metagenomic processing was performed, as the species-level OTUs were pre-defined. CLR abundance data (Table 2) input into the DENIM pipeline, yielded 5 distinct directed interaction networks. The reconstructed interaction network demonstrated a non-random, hierarchically organized topology, where species ranked by degree centrality, positioned highly connected taxa centrally and less connected ones peripherally across all axes. This spatial arrangement enabled clear visualization of interaction cascades: positive and negative symmetric interactions (+/+, −/−) clustered in quadrants I and III, while asymmetric polarity cascades (+/−, −/+) appeared in quadrants II and IV, facilitating comprehensive assessment of interaction dynamics across the network. The cumulative incoming interaction strength of a species closely mirrored species abundance trajectories consistently across replicates which can be seen in CLR abundance plots (Fig 1), further corroborated by quadrant-specific network analyses. Together, the network reveals an emergent ecological principle called polarity-driven succession, in which microbial community restructuring is governed by the balance of positive and negative interactions, with dominant taxa possessing more supportive positive links with magnitude over negative links, while suppressed species endure greater antagonistic pressures. This polarity-based framework offers a unified proximate explanation for abundance shifts and directional changes in microbial community structure during ecological succession and identifying the emerging interaction plasticity over the time.

**Table 1:**
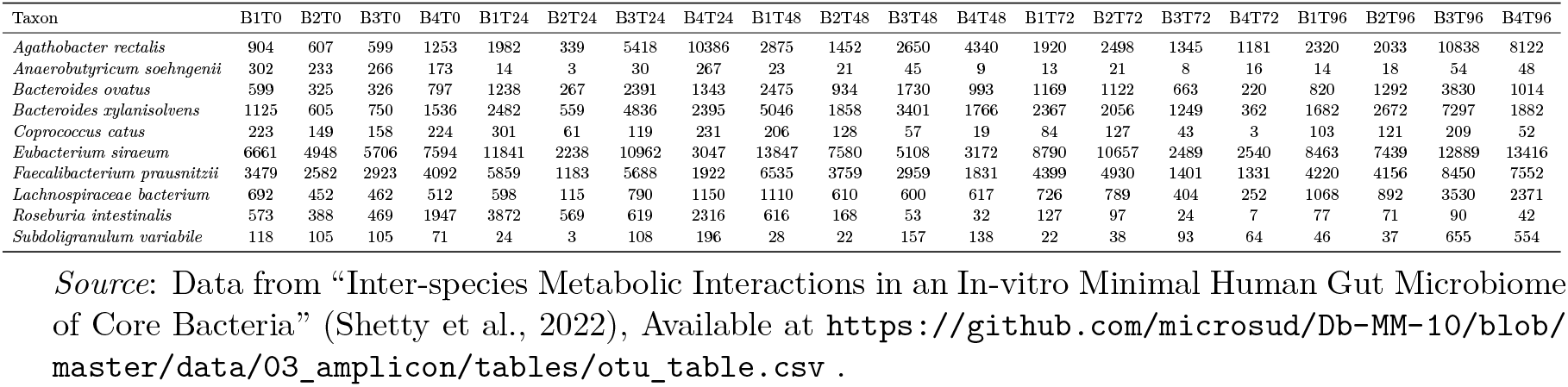
Abundances of microbial taxa across replicates.

**Table 2:**
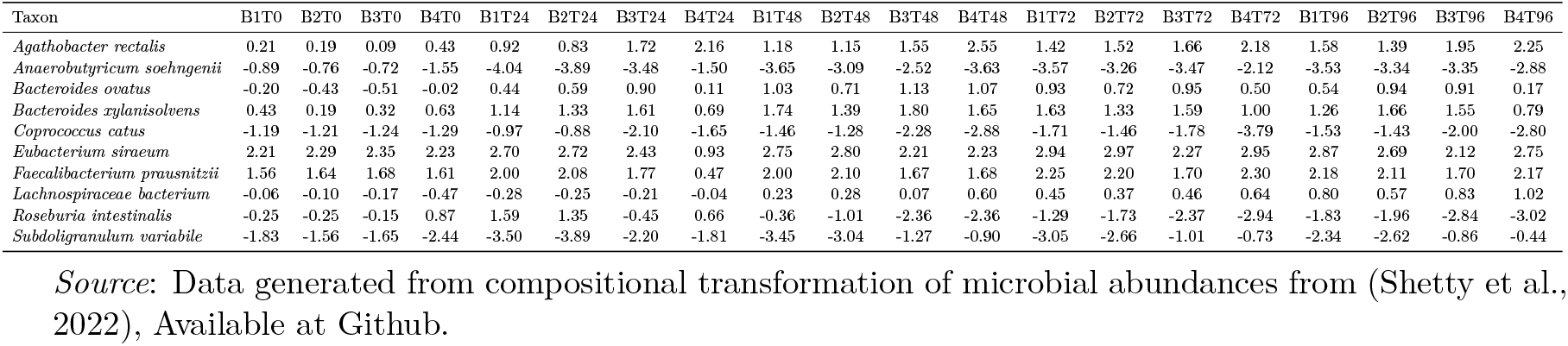
CLR-normalized abundances of microbial taxa across replicates.

**Table 3:** Mean CLR-normalized abundances of microbial taxa across replicates.

| Taxon | T0 | T24 | T48 | T72 | T96 |
| --- | --- | --- | --- | --- | --- |
| *Agathobacter rectalis* | 0.23172264598 | 1.40640406615 | 1.60673929504 | 1.69658282278 | 1.79136563858 |
| *Anaerobutyricum soehngenii* | -0.97969293658 | -3.22795648659 | -3.22289719845 | -3.10343972843 | -3.27616476084 |
| *Bacteroides ovatus* | -0.29254656998 | 0.51317191773 | 0.98364673755 | 0.77547840281 | 0.63781402565 |
| *Bacteroides xylanisolvans* | 0.39268794310 | 1.19250238895 | 1.64660292505 | 1.38609449630 | 1.31483927269 |
| *Coproccoccus catus* | -1.23291648956 | -1.39960406804 | -1.97702395871 | -2.18511117079 | -1.94254392099 |
| *Eubacterium siraeum* | 2.26953144699 | 2.19470561327 | 2.49857016141 | 2.78490922811 | 2.60800258461 |
| *Faecalibacterium prausnitzii* | 1.62273737955 | 1.58020925996 | 1.86164188804 | 2.11389829507 | 2.03927874977 |
| *Lachnospiraceae bacterium* | -0.19746686724 | -0.19496713889 | 0.29297010401 | 0.47847483208 | 0.80321404994 |
| *Roseburia intestinalis* | 0.05487842741 | 0.78579184730 | -1.52306343882 | -2.08309973212 | -2.41257715021 |
| *Subdoligranulum variable* | -1.86893497968 | -2.85025739983 | -2.16718651513 | -1.86378744581 | -1.56322848920 |

*Source:* Data generated from averaging the replicates of CLR-transformed abundance across samples.

**Figure 1:**
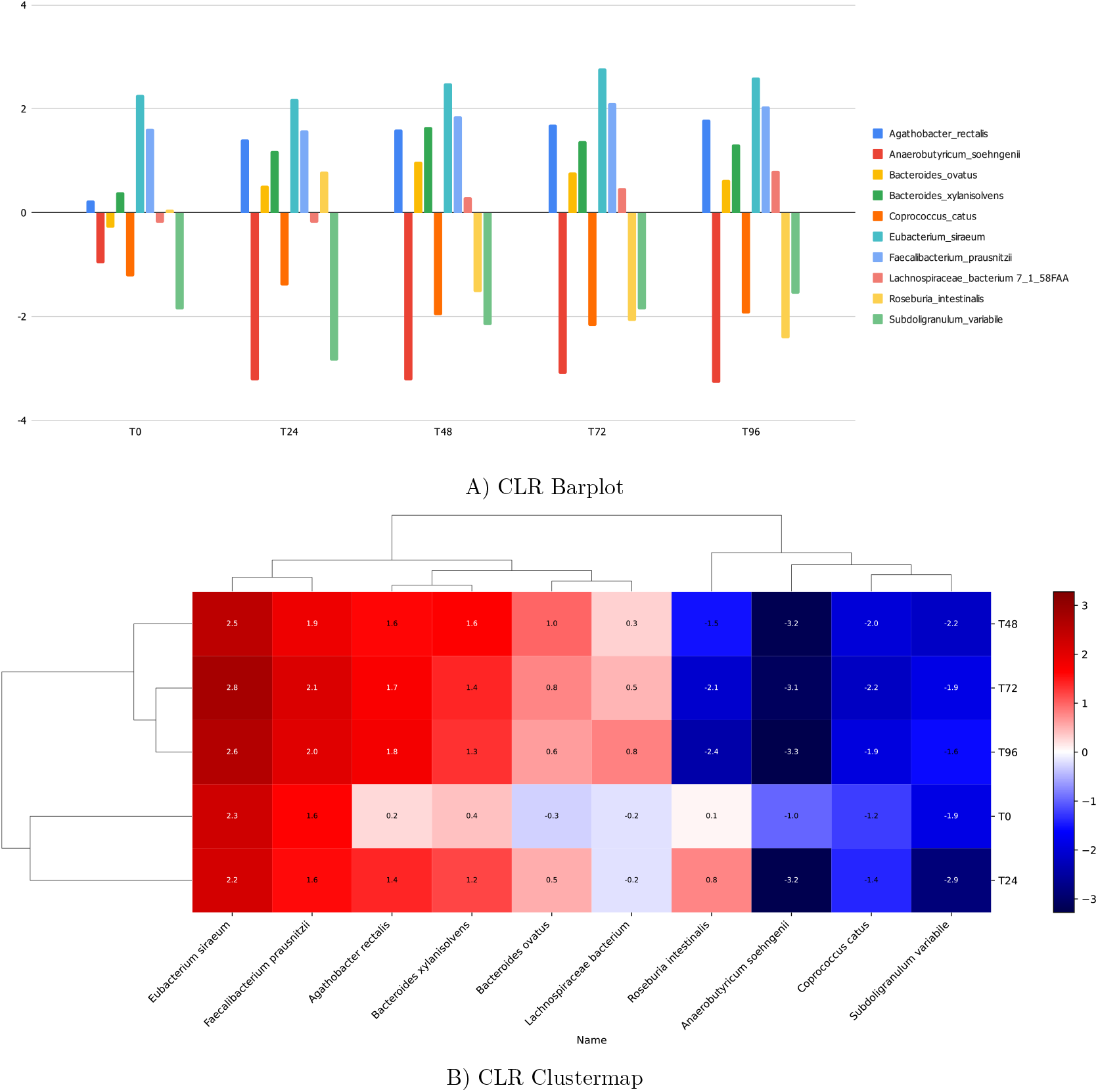
CLR-normalized mean Barplot and Clustermap of microbial taxa across replicates.

#### Box 1

##### Direction Assignment Proof

Given that the 16s based abundance data are inherently compositional, the centered log-ratio (CLR) transformation was applied (Gloor et al., 2017) to correct for compositional artifact. For each sample, the CLR value for species *i* was calculated as

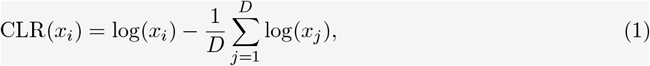

where *x*_*i*_ is the raw abundance of species *i*, and *D* = 10 is the total number of species. This transformation produces a matrix of CLR-transformed abundances suitable for downstream regression and correlation analyses. To identify statistically significant microbial associations, co-occurrence between species pairs was first examined. Analyses were restricted to samples in which both species exhibited non-zero CLR values to avoid zero-inflation biases. Pairs with fewer than three such samples (*n <* 3) were excluded.

Directionality of interactions between species pairs was inferred using a slope asymmetry heuristic based on regression. Given CLR abundance vectors х= (*x*_1_, *x*_2_, …, *x*_*n*_) and **y** = (*y*_1_, *y*_2_, …, *y*_*n*_), where *n ≥* 3, the following summary statistics were calculated:

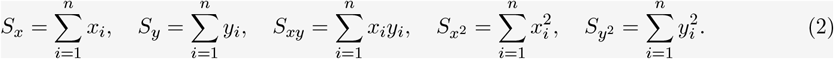

From these, the covariance and variances are defined as

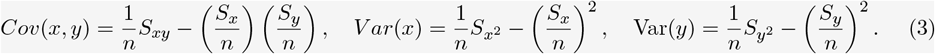

Linear regression slopes were then calculated as

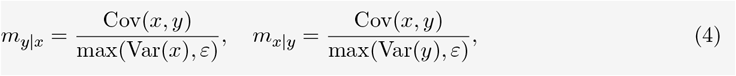

where *ε* is a small constant introduced to avoid division by zero.

A directional edge from *x* to *y* was assigned if the magnitude of one slope was at least 1.5 times greater than the other, that is,

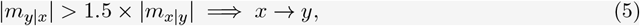

or vice versa. If neither direction exceeded this threshold, the pair was considered undirected and excluded. This approach captures asymmetries in ecological influence between taxa using steady-state abundance data.

#### Box 2

##### Sign Assignment Proof

After directionality was established, interaction types were classified via local perturbation analysis. For each directed pair (e.g., *x → y*), small perturbations were applied to the source species *x*, and the resultant changes in the target species *y* were evaluated. Perturbation magnitude at sample *i* was defined as

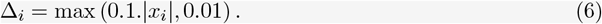

This means that for a given input value *x*_*i*_, the perturbation is set to either 10% of its absolute value or a fixed minimum of 0.01, whichever is greater. This adaptive approach ensures that the perturbation is proportional for larger values while maintaining a meaningful minimum effect for very small or near-zero values, preventing numerical instability or negligible changes during sensitivity analysis.

The perturbed values for each sample *i* are defined as

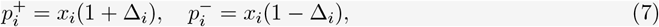

where Δ_*i*_ is the adaptive perturbation magnitude. Here, 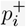 represents the value obtained by applying a positive perturbation to the original input *x*_*i*_, effectively increasing it by a proportion of Δ_*i*_. Conversely, 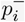 represents the negatively perturbed value, decreasing *x*_*i*_ by the same proportion. This symmetric perturbation around each input allows for evaluating the directional sensitivity of the system response.

To analyze the influence of an individual input feature *x*_*i*_ on the response variable *y*, directional perturbations are applied around the original feature value. Specifically, a *positive perturbation* 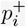 and a *negative perturbation* 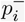 are introduced, and the corresponding estimated changes in the response are defined as:

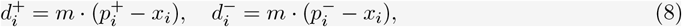

In this, *m* represents the estimated slope, which quantifies the local sensitivity of *y* with respect to *x*_*i*_. This slope captures how changes in the input feature influence the output in a linear approximation. The *sign* of each directional change indicates the qualitative effect of the perturbation on the target variable. A positive value suggests an increase in *y*, a negative value indicates a decrease, and a zero value implies no effect: sign 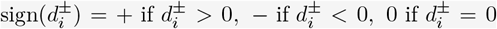. These sign-based classifications allow for the categorization of perturbation effects as positive, negative, or neutral effect on the target variable.

Let 𝒩 be the set of all combined perturbation-effect notations observed over all replicates, where each notation *n* ∈ 𝒩has the form: *n* = *σ/ϵ* with *σ* ∈ {+, *−*}denoting the **perturbation direction** (positive or negative) and *ϵ* ∈ {+, *−*} denoting the **sign of the observed effect** on the target variable. Each notation *n* has an associated count *c*_*n*_ ∈ ℕ, representing the frequency of observing that combined perturbation-effect pattern: *c*_*n*_ = # { *i*: Frequency of observed notations *i* = *n* }

The final interaction notation, sign, is defined as the notation with the highest observed count: sign = max_*n*∈𝒩_ *c*_*n*_. If no notations are observed (𝒩= *∅*), no sign is assigned.

## 3 RESULTS AND DISCUSSION

### 3.1 Temporal Shifts in Abundance Dynamics

CLR-normalized barplot and clustermap (Fig 1) illustrated temporally resolved microbial abundance dynamics, revealing distinct compositional shifts across the experimental timepoints. Several taxa, including *Agathobacter rectalis, Bacteroides xylanisolvens, Bacteroides ovatus*, and *Lachnospiraceae bacterium*, exhibited a consistent increase in CLR values, suggesting successful adaptation and a potential ecological advantage under the imposed conditions. Conversely, *Anaerobutyricum soehngenii, Roseburia intestinalis*, and *Coprococcus catus* showed marked declines, indicative of environmental sensitivity or suppression by competitive interactions. *Eubacterium siraeum* and *Faecalibacterium prausnitzii* maintained stable, high CLR values, positioning them as resilient core members of the microbiota.

### 3.2 Interspecies Dependencies and Ecological Roles

To probe interspecies ecological dependencies, all 45 possible pairwise interactions among the 10 dominant taxa were assessed across five timepoints (T0, T24, T48, T72, T96) using permutation-based association testing, where the DENIM pipeline was applied to infer the interactions. On average, 37 interactions per timepoint were seen, with interaction counts of 40, 36, 37, 37, and 36 were shortlisted respectively. Eco-Cartesian plots (Fig 2) of these interactions highlighted structured, directional associations that evolved over time, capturing the abundance based dependence. Higher-order microbial interactions were further examined through Directed Acyclic Graphs (DAGs), capturing conditional dependencies among taxa, though multiple DAGs were found only 9 were represented distinctly, which has species > 2 and spanning time-points > 2. Among the nine DAGs, five persisted consistently from T48 to T96 with identical signs across time, which indicates and persistent interaction establishments, which also explains the origin of interaction plasticity between species pairs (Solowiej-Wedderburn et al., 2025). These persistent plastic interaction motifs were predominantly initiated by *Lachnospiraceae bacterium*, suggesting its potential role as a keystone or ecological initiator. Intermediate nodes within these motifs frequently included *Agathobacter rectalis, Anaerobutyricum soehngenii, Roseburia intestinalis*, and *Eubacterium siraeum*, while *Subdoligranulum variabile* was often observed as a terminal responder, reflecting its downstream positioning in the network.

**Figure 2:**
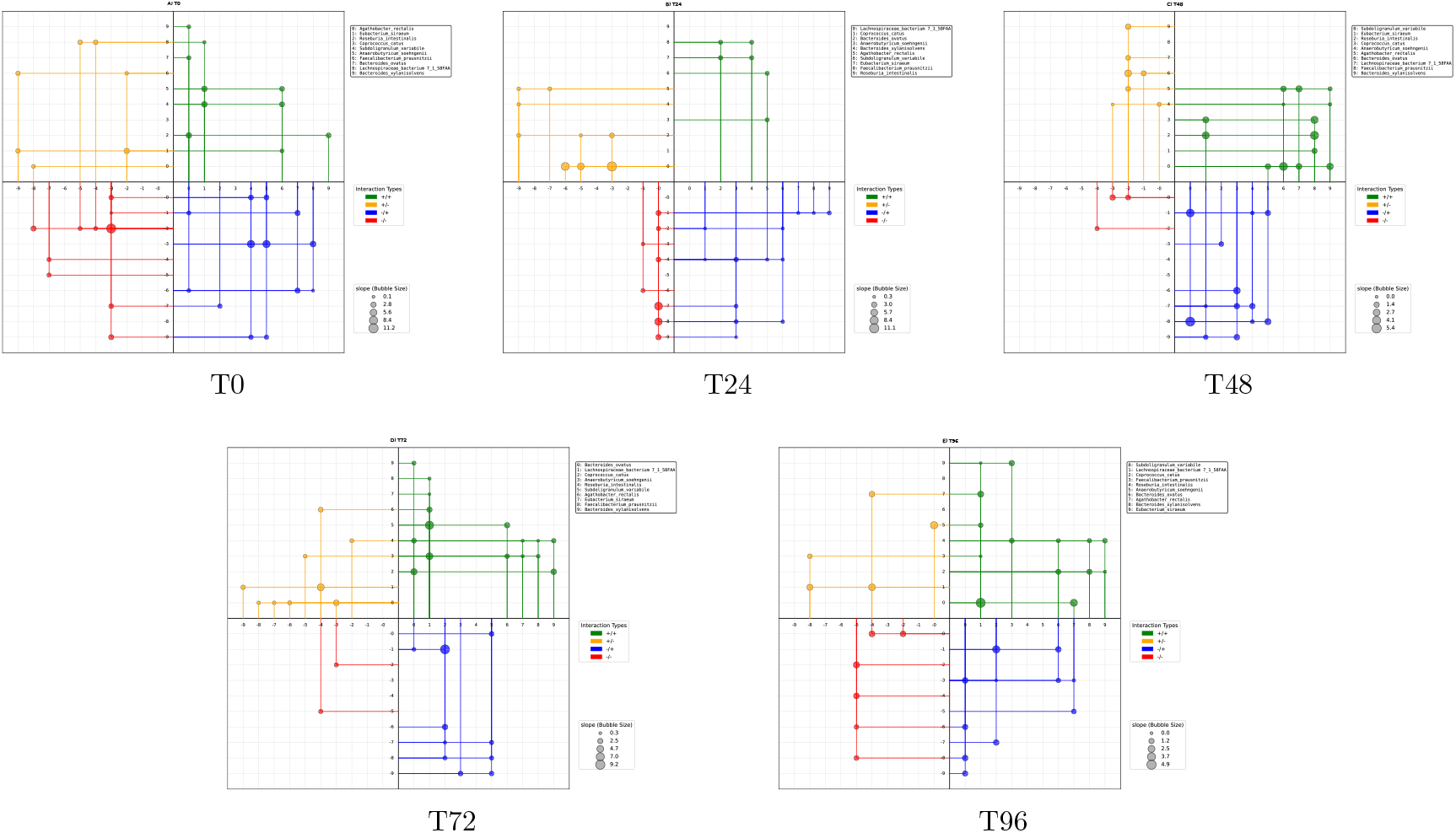
Directed, signed Eco-Cartesian interaction network inferred from centered log-ratio (CLR) normalized abundance data. Nodes placed on all the axis, that represent microbial species ordered by degree centrality, positioned according to a custom quadrant-based layout that separates outgoing (influencing) from one axis to incoming (receiving) interactions on next axis in counter-clockwise direction where (quadrant: Q) Q1: +/+ is +*x* → +*y*, Q2: +/− is +*y* → *− x*, Q3: −/− is *−x* → *−y* and Q4: −/+ is *− y →* +*x*. Edge direction indicates the inferred ecological influence from one taxon to another; edge color denotes interaction type; and edge bubbles corresponds to interaction strength (slope magnitude). This layout facilitates intuitive interpretation of species-level roles, highlighting ecological influencers (+ve influencers from I and II quadrants, −ve influencers from III and IV quadrants), responders (+ve responders from I and IV quadrants, −ve responders from II and III quadrants), and taxa under net antagonistic or cooperative pressure. The visualization mitigates the hairball effect and allows direct comparison of network dynamics with abundance trends within a sample and across time points at T0, T24, T48, T72, and T96 hours.

### 3.3 Network Topology and Visualization

The reconstructed Eco-Cartesian interaction network demonstrated a non-random, hierarchically organized topology. Species were positioned by degree centrality with highly connected taxa centrally located and less connected species arranged peripherally across all axes. This spatial arrangement enabled clear visualization of interaction cascades: positive and negative symmetric interactions (+/+, −/−) clustered in quadrants I and III, while asymmetric polarity cascades (+/−, −/+) appeared in quadrants II and IV, facilitating comprehensive assessment of interaction dynamics across the network. Temporal shifts in the cumulative incoming interaction strength closely mirrored species abundance trajectories consistently across replicates which can be seen in CLR abundance plots (Fig 1), further corroborated by quadrant-specific network analyses. Together, these results reveal an emergent ecological principle in which microbial community restructuring is governed by the balance of positive and negative interactions, with dominant taxa possessing more supportive positive links with magnitude over negative links, while suppressed species endure greater antagonistic pressures. This polarity-based framework offers a unified explanation for abundance shifts and directional changes in microbial community architecture during ecological succession. Additionally the Eco-Cartesian layout that separates outgoing (influencing) from one axis to incoming (receiving) interactions on next axis for each quadrant in counter-clockwise direction where (Quadrant: Q) Q1: +/+ is +*x* → +*y*, Q2: +/− is +*y* → *− x*, Q3: −/− is *−x* → *− y* and Q4: −/+ is *−y* →+*x*. Though unconventional these counter-clockwise direction across all quadrant and heuristic axes flips in-between the quadrants facilitates intuitive interpretation of species-level interactions in cyclical manner, highlighting ecological influencers (+ve influencers from I and II quadrants, −ve influencers from III and IV quadrants), responders (+ve responders from I and IV quadrants, −ve responders from II and III quadrants), and taxa under net antagonistic or cooperative pressure. This visualization strategy mitigates the hairball effect and allows direct comparison of network dynamics with abundance trends within a sample.

### 3.4 Methodological Advancements and Comparison with Existing Approaches

Decreased microbial diversity has been associated with health-compromised hosts, which may limit their plasticity and ability to respond to environmental change (Stevick et al., 2021), thus microbiota plasticity is defined as variability in the structure and composition of the microbiota (Grembi et al., 2020). Understanding the ecological interaction’s plasticity that underpin microbial community structure is central to deciphering their functional dynamics and stability. Microbial ecosystems exhibit a wide range of interspecies relationships, from mutualism and syntrophy to competition and antagonism, which collectively shape their composition and resilience. However, inferring these interactions remains challenging due to the compositional nature of sequencing data, high dimensionality, and inherent sparsity in microbiome datasets. Traditional approaches, such as taxonomic bar plots or correlation-based networks, offer only coarse, de-scriptive insights and fail to capture directionality, causality, or mechanistic underpinnings of microbial interactions (Srinivasan et al., 2024; Röttjers and Faust, 2018; Layeghifard et al., 2017), which could be resolved with minor caveats. A major limitation of many existing methods is their dependence on undirected co-occurrence or correlation matrices, which are prone to conflating direct and indirect associations, but still widely adapted to represent microbial associations, though correlation is considered as a proxy for interaction (Fisher and Mehta, 2014). Furthermore, such methods typically do not account for the statistical distortions introduced by the compositionality of relative abundance data, often leading to spurious inferences (Gloor et al., 2017). Although tools like SparCC and SPIEC-EASI address some of these constraints, they generally produce undirected graphs and are unable to infer interaction sign or strength, which limits their utility in modeling dynamic community behavior (Faust and Raes, 2012). To overcome these barriers, this heuristic network inference pipeline was developed, which also represent associations supported with directed, signed, and weighted microbial interaction networks from cross-sectional compositional datasets (Xiao et al., 2017). Without relying on time-series data or predefined mechanistic models, this method leverages asymmetric slope estimation and dual-value perturbation heuristics to infer the directionality and polarity of interactions. The approach thus enables a shift from static, descriptive microbiome profiling to mechanistic ecological interpretation, allowing microbial taxa to be categorized based on their functional roles such as ecological influencers, responders, or structural hubs.

**Figure 3:**
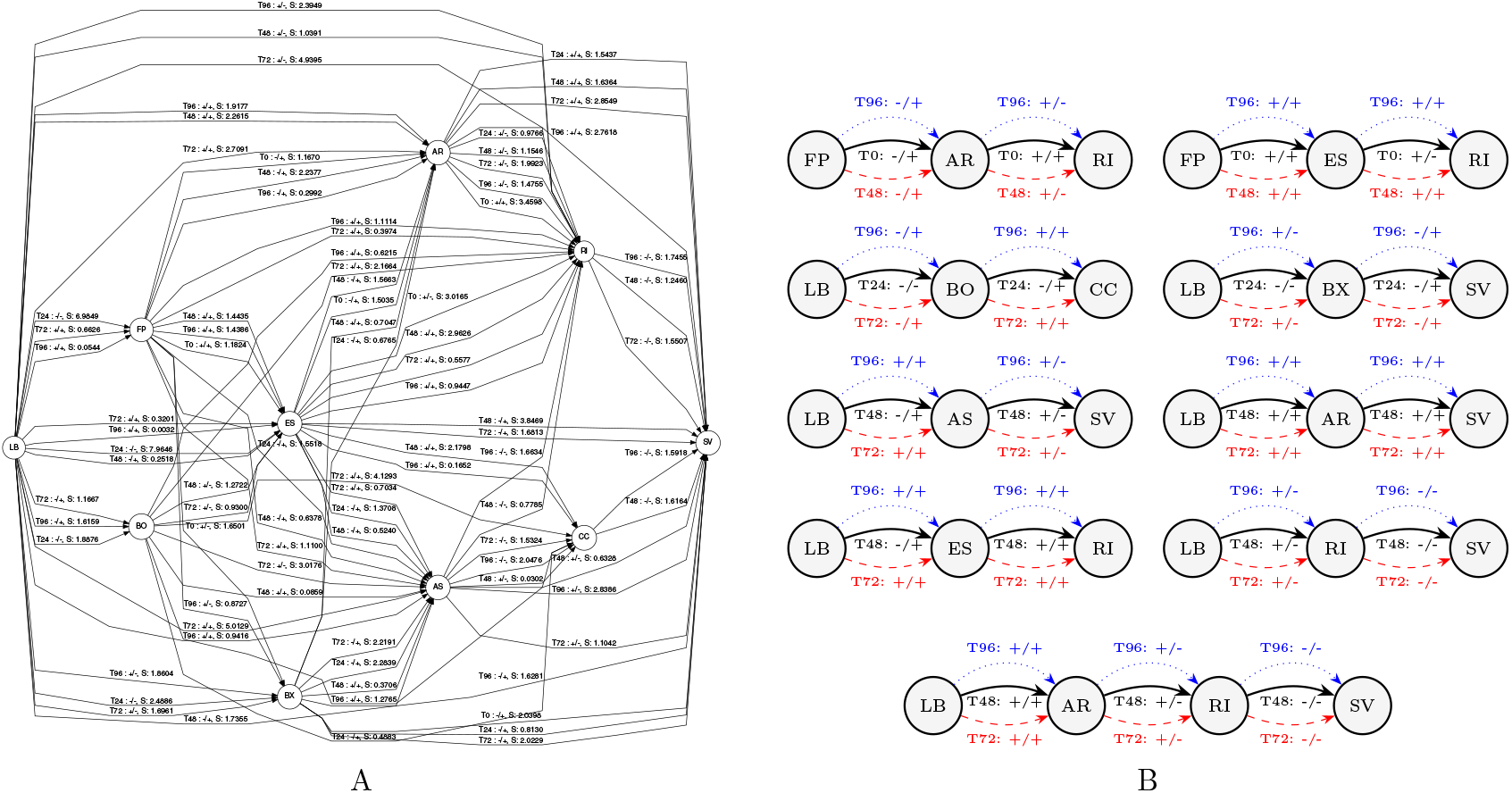
The Directed Acyclic Graph (DAG) plots A & B illustrate the inferred directional interactions among microbial taxa that persist across multiple timepoints, revealing hierarchical ecological dependencies within the community. Each node represents a microbial species, and arrows indicate the direction of influence along with slope (S), capturing the potential flow of ecological impact from one taxon to another. These DAGs highlight conditionally independent, persistent high-order interaction plasticity, emphasizing taxa that consistently act as initiators or responders over time. For clarity in visualization, each taxon is represented by a two-letter abbreviation: AR for *Agathobacter rectalis*, AS for *Anaerobutyricum soehngenii*, BO for *Bacteroides ovatus*, BX for *Bacteroides xylanisolvens*, CC for *Coprococcus catus*, ES for *Eubacterium siraeum*, FP for *Faecalibacterium prausnitzii*, LB for *Lachnospiraceae bacterium*, RI for *Roseburia intestinalis*, and SV for *Subdoligranulum variabile*.

One of the key advantages of this approach is its capacity to identify critical taxa based on network topology. By employing degree centrality metrics, where the nodes with extensive connectivity and control over interaction flow are prioritized. In a microbiome context, such nodes may serve as ecological keystones or hubs, influencing the broader community through direct and indirect interactions (Layeghifard et al., 2017; Oña et al., 2025) and the conditional persistent DAGs facilitates the identification of established persistent interactions and high degree of plasticity depending on the environment (Solowiej-Wedderburn et al., 2025). A distinguishing feature of this work is the introduction of a custom visualization technique tailored for ecological networks. Unlike conventional network diagrams that often result in visually dense and uninterpretable hairballs due to undirected, overlapping edges (Röttjers and Faust, 2018), the proposed layout separates microbial abundance dependent interactions into directional quadrants that distinguish between outgoing and incoming interactions. This quadrant-based visualization enhances interpretability by directly linking topological attributes with ecological function. For instance, taxa with high incoming negative influence and magnitude often correspond with reduced relative abundance and vice versa, while those with strong outgoing positive influence and magnitude may drive community dynamics and vice versa as important ecological drivers. The design minimizes edge clutter and retains topological clarity even as network size increases, making it suitable for high-complexity datasets. The inferred interaction networks also enable deeper ecological interpretation. Taxa with high cumulative outgoing influence can be viewed as potential keystone species (Oña et al., 2025), while others consistently positioned as recipients of negative interactions may represent suppressed or competitively disadvantaged members. Moreover, motif analysis within these networks reveals ecologically relevant structures such as cascades, control modules, and signal integration patterns providing mechanistic insights into community level functions (Ganter et al., 2014). The using of publicly available synthetic dataset, to identify key driver taxa, conditionally persistent DAG motifs, and temporal rewiring of microbial influence patterns shows that, Although the dataset was not generated specifically for this method, the consistent emergence of interpretable ecological structures such as the initiator role of *Lachnospiraceae bacterium* supports the validity of the approach.

### 3.5 Practical Implications

Understanding individual microbes as components of a complex biological system is essential. To reverse-engineer the effects of a microbial community, it is necessary to study both the individual members and the interactions between them. By investigating microbial composition and interrelationships, we can gain deeper insight into the direct and indirect roles of specific taxa, anticipate community responses to environmental perturbations, and ultimately inform the rational design and engineering of microbial consortia for beneficial applications (Kodera et al., 2022; Bernabe et al., 2018), which can be intuitively supported by DENIM. This network-based framework has practical implications for microbial cultivation and synthetic community design. By identifying potential mutualists or antagonists, it facilitates the strategic co-culturing of fastidious or previously uncultured microbes. Such insights are especially valuable in biotechnological and clinical contexts, where controlled manipulation of microbial communities is essential (Lv et al., 2019). Additionally, the method is well-suited for integration with the rapidly expanding repository of publicly available microbiome datasets. Its reliance solely on cross-sectional relative abundance data allows retrospective application across thousands of studies. This opens paths for resource efficient research, particularly in settings with limited access to experimental infrastructure, and promotes equitable participation in global microbiome science (Kelliher et al., 2025; Jurburg et al., 2024). When used in combination with advanced visualization tools such as Cytoscape and Hive plots, the interpretability and accessibility of these ecological networks are further enhanced (Krzywinski et al., 2012). As microbiome datasets continue to grow in scale and complexity, the need for methods that balance computational rigor with ecological relevance becomes increasingly important. The approach outlined here represents a scalable, data efficient, and mechanistically informative solution to microbial interaction inference, paving the way for more predictive and hypothesis driven microbiome research.

In summary, the developed network inference method provides a principled framework for mapping microbial ecological interactions from cross-sectional compositional data. It addresses key methodological challenges and enables a more nuanced understanding of microbial community structure, ecological roles, and potential functional dynamics (Xiao et al., 2017).

## 4 CONCLUSIONS

The DENIM pipeline proposed in this work overcomes the directionality prediction and interaction classification between species by inferring directed, signed, and weighted ecological interaction networks from cross-sectional, compositional 16S rRNA sequencing data. By addressing core limitations of traditional correlation-based approaches including lack of directionality, ecological polarity, and compositional bias this method enables a more mechanistically grounded interpretation of microbial community interactions. The framework operates under the assumption of quasi-steady-state conditions within individual timepoints, allowing for meaningful ecological inference in the absence of dense time-series data or predefined dynamic models. It leverages asymmetric slope estimation and adaptive perturbation heuristics to infer interaction direction, sign, and strength while accounting for the constraints of compositional data. Validation on a publicly available, temporally resolved synthetic dataset demonstrated the framework’s capacity to recover consistent and ecologically plausible interaction patterns, including conditionally persistent network motifs and interpretable topological structures. These findings support the method’s robustness and generalizability and across datasets that are not originally designed for network inference. The biologically optimized quadrant-based visualization strategy further enhanced the interpretability of the resulting networks, facilitating the identification of functionally distinct ecological roles without introducing visual complexity typical of traditional network layouts. Overall, this approach provides a scalable, data-efficient, and biologically interpretable tool for microbial interaction inference. Its compatibility with widely available cross-sectional datasets offers new opportunities for predictive microbiome research, mechanistic hypothesis generation, and the design of targeted experimental or synthetic community interventions.

## Supporting information

OTU Abundance table

CLR Normalised OTU table

Mean CLR table of Replicate averaged OTU table

Overall edge interaction table

Filtered network table

## Abbreviations

DAGs: Directed Acyclic Graphs
CLR: Centered Log-ratio Normalization

## DATA AVAILABILITY

All the datasets analyzed in this study are either publicly available or kindly provided by the original authors. Other data that supports the findings of this study are available from the corresponding author on reasonable requests.

## ACKNOWLEDGMENTS

The author acknowledges the creators of the publicly available dataset from the study “Inter-species Metabolic Interactions in an In-vitro Minimal Human Gut Microbiome of Core Bacteria” (Shetty et al., 2022).

## SUPPORTING INFORMATION

Additional supporting information may be found in the online version of the article at the publisher’s website.

